# Investigating the biosynthesis and roles of the auxin phenylacetic acid during *Pseudomonas syringae-Arabidopsis thaliana* pathogenesis

**DOI:** 10.1101/2024.04.04.587729

**Authors:** Chia-Yun Lee, Christopher P. Harper, Soon Goo Lee, Yunci Qi, Taylor Clay, Yuki Aoi, Joseph M. Jez, Hiroyuki Kasahara, Joshua A. V. Blodgett, Barbara N. Kunkel

## Abstract

Several plant-associated microbes synthesize the auxinic plant growth regulator phenylacetic acid (PAA) in culture; however, the role of PAA in plant-pathogen interactions is not well understood. In this study, we investigate the role of PAA during interactions between the phytopathogenic bacterium *Pseudomonas syringae* strain *Pto*DC3000 (*Pto*DC3000) and the model plant host, *Arabidopsis thaliana*. Previous work demonstrated that indole-3-acetaldehyde dehydrogenase A (AldA) of *Pto*DC3000 converts indole-3-acetaldehyde (IAAld) to the auxin indole-3-acetic acid (IAA). Here, we further demonstrate the biochemical versatility of AldA, as it can use both IAAld and phenylacetaldehyde as substrates to produce IAA and PAA, respectively. We also show that during infection AldA-dependent synthesis of either IAA or PAA by *Pto*DC3000 does not contribute significantly to the increase in auxin levels in *A. thaliana* leaves. Using available *arogenate dehydratase* (*adt*) mutant lines of *A. thaliana* compromised for PAA synthesis, we observed that a reduction in PAA-Asp and PAA-Glu is correlated with elevated levels of IAA and increased susceptibility. These results provide evidence that PAA/IAA homeostasis in *A. thaliana* influences the outcome of plant-microbial interactions.

## 1 Introduction

Auxins are a class of phytohormones that regulate various aspects of plant growth and development; auxins also participate in many plant-microbe interactions (Kunkel and Harper, 2018; Spaepen and Vanderleyden, 2011). For example, indole-3-acetic acid (IAA), one of the best-known auxins, promotes pathogen colonization and growth by suppressing host defense responses (Chen et al., 2007; Djami-Tchatchou et al., 2020; McClerklin et al., 2018; Wang et al., 2007). Additionally, IAA serves as an environmental signal that can modulate bacterial gene expression related to antimicrobial tolerance, stress responses, and pathogenesis (Djami-Tchatchou et al., 2022; Djami-Tchatchou et al., 2020; Spaepen et al., 2007; Van Puyvelde et al., 2011; Yuan et al., 2008).

Phenylacetic acid (PAA) is another form of natural auxin that can be synthesized by both plants (Aoi et al., 2020b; Cook et al., 2016; Dai et al., 2013; Perez et al., 2023; Sugawara et al., 2015) and microorganisms (Akram et al., 2016; Bartz et al., 2013; Slininger et al., 2004; Sopheareth et al., 2013; Spaepen et al., 2007). PAA acts either as a carbon and energy source for microbes or as a signaling molecule that induces chemotaxis, catabolism, and modulation of virulence gene expression (Bhuiyan et al., 2016; Wang et al., 2013); however, the biological role of PAA in plant-associated microbes is not well understood.

Previously, we demonstrated that the plant pathogen *Pseudomonas syringae* strain *Pto*DC3000 produces IAA via indole-3-acetaldehyde dehydrogenase A (AldA), which catalyzes the NAD-dependent formation of IAA from indole-3-acetaldehyde (IAAld) (McClerklin et al., 2018). Because there are several known parallels in IAA and PAA metabolism (Cook et al., 2016; Dai et al., 2013; Somers et al., 2005; Sugawara et al., 2015; Tao et al., 2008), we speculated that AldA might have a role in PAA biosynthesis via a phenylacetaldehyde (PAAld) intermediate.

In this study, we explore the AldA-dependent auxin synthesis pathway in *Pto*DC3000 and its role in pathogenesis on *Arabidopsis thaliana*. We first demonstrate that *Pto*DC3000 AldA can convert PAAld to PAA. We also show that AldA-dependent auxin synthesis does not significantly contribute to the increase in auxin levels within infected plant tissues, suggesting pathogen infection stimulates production of auxin by the host. To further investigate the role of PAA during *Pto*DC3000 pathogenesis, we observe that a reduction in PAA-Asp and PAA-Glu correlates with elevated IAA and increased susceptibility to *Pto*DC3000. Thus, while *Pto*DC3000 auxin synthesis does not apparently contribute to the increased auxin in infected plant tissue, *Pto*DC3000 infection perturbs PAA/IAA homeostasis, which influences the outcome of the interaction.

## 2 Materials and methods

### 2.1 Bacterial strains and plasmids

The bacterial strains and plasmids used in this study are summarized in Table S1. *P. syringae* strain *Pto*DC3000 (*Pto*DC3000) wild-type (Cuppels Diane, 1986) and mutant strains were grown on Nutrient Yeast Glycerol (NYG) medium (Daniels et al., 1988) and Hoitkin-Sinden medium supplemented with 10 mM citrate (HSC) (Sreedharan et al., 2006) at 28-30°C. *Escherichia coli* was grown on Luria Broth (LB) medium at 37°C. Antibiotics used for selection include: rifampicin (Rif, 80 µg/mL), kanamycin (Km, 25 µg/mL), tetracycline (Tet, 16 µg/mL), spectinomycin (Spec, 100 µg/mL), and chloramphenicol (Cm, 20 µg/mL).

### 2.2 Construction of mutant *Pto*DC3000 strains

The primers used in this study are listed in Table S2. A plasmid for generating the *aldA*::omega (Ω) deletion mutant was constructed from the suicide vector pJP5603 (Penfold and Pemberton, 1992). The Ω fragment containing Spec resistance was amplified from purified pHP45 plasmid DNA (Prentki and Krisch, 1984) using the primers omega_frag_F and omega_frag_R. The vector pJP5603 sequence was linearized by PCR using the primers pJP5603_F and pJP5603_R, followed by DpnI digestion of the template plasmid. Fragments of approximately 1 kb corresponding to genomic regions upstream and downstream of the *aldA* (*PSPTO_0092*) gene were amplified from *Pto*DC3000 genomic DNA by PCR using primers ald_up_F and ald_up_R, and ald_down_up and ald_down_R, respectively. The fragments were assembled using NEB HiFi DNA Assembly Master Mix (Ipswich) to create plasmid pJP5603-*aldA*::Ω, transformed into *E. coli* DH5α λpir cells (Miller and Mekalanos, 1988) and plated onto LB medium containing Spec. The plasmid was sequenced to confirm that the assembly occurred correctly and that no mutations were inadvertently introduced.

pJP5603-*aldA*::Ω was conjugated into *Pto*DC3000 using the helper strain MM294A (pRK2013) (Finan et al., 1986) to create a single crossover plasmid insertion, which was confirmed by PCR genotyping. The single crossover strain was grown in NYG Spec and subcultured for 13 days to obtain a mutant in which a double crossover event occurred. Approximately 8000 colonies were screened for loss of the Km-resistance cassette by replica plating until a Spec-resistant, Km-sensitive mutant was obtained. The resulting strain was genotyped by PCR using primers omega_out_up and omega_out_down, to verify that the wild-type *aldA* gene had been replaced by the omega fragment. The *aldA*::Ω *aldB*::pJP5603-Km double mutant was generated by introducing the pJP5603-2673int insertional disruption plasmid (McClerklin et al., 2018) into the *aldA*::Ω mutant by triparental mating and selecting for Km-and Spec-resistant colonies. Disruption of *aldB* (*PSPTO_2673)* by pJP5603 was confirmed by PCR using primers M13F and 2673SeqF.

### 2.3 Feeding *P. syringae* with PAAld

Wild-type *Pto*DC3000 and mutant strains were grown in NYG medium without antibiotics until they entered the exponential phase of growth. These cultures were used to inoculate HSC medium at a density of ∼1×10^7^ CFU mL^-1^ and incubated with shaking for 48 hours (hrs) at 28°C. The culture medium was supplemented with 25 µM PAAld (Sigma Aldrich) in 0.24% ethanol (EtOH), 100 µM phenylalanine (Sigma Aldrich) in 0.24% EtOH, IAAld-sodium bisulfite (Sigma Aldrich) in 0.24% EtOH, or 0.24% EtOH (mock) as indicated. Samples (1 mL) were taken 46-48 hrs after addition, pelleted by centrifugation, and the resulting supernatants filtered with a 0.2-micron filter and stored at −80°C until quantification. Growth of cultures was monitored by reading the OD_600_ at regular intervals with a spectrophotometer. Conditioned HSC was made by growing *Pto*DC3000 in HSC for 48 hrs followed by removing the bacterial cells by centrifugation and filtering the supernatant through a 0.2-micron filter. PAAld (final concentration: 25 µM) was added to the conditioned HSC. After 48-hr incubation at 28°C, medium was collected and PAA levels quantified.

### 2.4 Quantification of PAA production in culture

Metabolite analysis was performed using a Phenomenex Luna Omega polar C18 column (50 × 2.1 mm, 3 μm pore size) installed on an Agilent 1260 Infinity HPLC connected to an Agilent 6420 Triple-Quad mass spectrometer. Metabolites were separated using the following chromatography conditions: T = 0, 0% B; T = 2, 0% B; T = 3, 20% B; T = 8, 40% B, T = 10, 100% B; T = 12, 100% B; T = 14, 0% B; T = 16, 0% B; where buffer A was water + 0.1% formic acid and buffer B was acetonitrile + 0.1% formic acid and the flow rate was 0.5 mL/min. The column was held at 20°C, and 8 μL of the sample was injected per run. For quantification, the mass spectrometer was set to multiple reaction monitoring in positive ion mode. The mass transitions for each metabolite were chosen using Agilent Optimizer. The mass transitions, fragmentation conditions, and retention times for each metabolite are listed in Table S3. For d5-Trp (CDN Isotopes Inc.) labeling experiments, the mass spectrometer was set to scan for m/z 100-400 in positive MS2 scan mode. The resulting data were analyzed offline with the Agilent MassHunter Quantitative Analysis and Qualitative Analysis software.

### 2.5 Protein expression and purification

AldA was expressed and purified using the pET28a-AldA construct as described (McClerklin et al., 2018). Hexahistidine-tagged AldA was further purified through a Superdex-200 26/60 size-exclusion column (SEC) (GE healthcare) in 25 mM HEPES, 100 mM NaCl, pH 7.5. Purified AldA was stored in SEC buffer containing 20% glycerol (25 mM HEPES, 100 mM NaCl, 20% (v/v) glycerol, pH 7.5) at −80°C. To determine the concentration of AldA for enzymatic analysis, the molar extinction coefficient (ε_280 nm_ = 68,410 M^-1^ cm^-1^) at A_280nm_ calculated using ProtParam (Gasteiger et al., 2005) was employed.

### 2.6 Aromatic aldehyde substrate screening and steady-state kinetic analysis

Enzymatic activity of AldA was measured by continuously monitoring the spectrophotometric absorbance changes from NAD to NADH (ε_340 nm_ = 6220 M^−1^ cm^−1^) at A_340nm_ using an EPOCH2 microplate spectrophotometer (BioTek). Substrate screening experiments were conducted at 25°C with 1 mM of NAD^+^ and 5 mM of each aldehyde (*i*.*e*., IAAld, PAAld, hydrocinnamaldehyde, and cinnamaldehyde) in the assay conditions of 100 mM Tris·HCl (pH 8.0) and 100 mM KCl. Steady-state kinetic parameters of AldA for PAAld were determined in the same condition with varied PAAld (0.01–2.5 mM) and fixed cofactor (NAD^+^; 5 mM) or with fixed PAAld (1 mM) and varied cofactor (NAD^+^; 0.05–2.5 mM). The resulting initial velocity data were fit to the Michaelis–Menten equation, *v* = (*k*_cat_[S])/(*K*_m_ + [S]), using Prism (GraphPad).

### 2.7 Computational docking

Molecular docking experiments for AldA were performed by AutoDock vina (Version 1.1.2) (Gasteiger et al., 2005; Trott and Olson, 2010) with standard protocols as previously described (McClerklin et al., 2018). Docking of IAAld and PAAld into the AldA active site used a 30 × 30 × 30 Å grid box with the level of exhaustiveness = 20. The x-ray crystal structure of the AldA• NAD^+^•IAA complex (PDB: 5IUW) was used as a template with fixed position of NAD^+^ (McClerklin et al., 2018). Docking of IAAld and PAAld yielded a calculated affinity of −5.9 to −4.2 kcal mol^−1^ and −5.3 to −3.3 kcal mol^−1^, respectively.

### 2.8 Plant material and growth conditions

All *A. thaliana* mutants and transgenic lines used in this study were in the Col-0 background. The *adt1 adt3 adt4 adt5 adt6* quintuple mutant (*adt1/3/4/5/6)* and *ADT4* and *ADT5* overexpression transgenic lines (ADT4 OE and ADT5 OE) have been previously described (Aoi et al., 2020b; Chen et al., 2016). Plants were grown on soil in a growth chamber with a short-day photoperiod (8 hrs light/16 hrs dark) at 21°C and 75% relative humidity, with a light intensity of ∼ 130 μEinsteins sec^-1^ m^-1^.

### 2.9 *P. syringae* inoculation and quantification of bacterial growth

*A. thaliana* plants were infected at approximately four weeks of age. For bacterial growth quantification, 10^5^ cells mL^-1^ were resuspended in 10 mM MgCl_2_ and injected into leaves using a 1-mL needleless syringe. Whole leaves were sampled at ∼2 hrs after inoculation (day 0) and 4 days-post-inoculation (dpi), weighed to determine leaf fresh mass, ground in 10 mM MgCl_2_ and then plated in serial dilutions on NYG media with rifampicin. Following incubation at 28°C for 48 hrs, colonies were counted to determine the number of bacteria in the leaves. Between four and eight leaves were sampled per treatment, depending on the experiment. For disease symptom observations, 10^6^ cells mL^-1^ were resuspended in 10 mM MgCl_2_ and infiltrated into leaves. Disease symptoms were photographed at 4 dpi.

### 2.10 Quantification of auxin and auxin-amino acid conjugates *in planta*

IAA, IAA-Asp, IAA-Glu, PAA, PAA-Asp, and PAA-Glu were extracted from freeze-dried plant material, purified and analyzed by an Agilent 6420 Triple Quad system (Agilent Technologies, Inc.) with a ZORBAX Eclipse XDB-C18 column (1.8 µm, 2.1 × 50 mm) as previously reported (Aoi et al., 2020c).

### 2.11 Statistical analysis

Datasets were statistically compared with the statistical analysis software GraphPad Prism 9.5.0 (GraphPad). Statistical tests included Student’s *t*-test or one-way analysis of variation (ANOVA) followed by the Tukey’s HSD test when appropriate. The confidence level of all analyses was set at 95%, and values with *p* < 0.05 were considered significant.

## 3 Results

### 3.1 *P. syringae* strain *Pto*DC3000 produces PAA in culture

Previous biochemical experiments carried out in *Azospirillum brasilense* suggested that PAA biosynthesis proceeds via the IAA biosynthetic enzyme indole pyruvate decarboxylase, by converting phenylpyruvate to PAAld (Somers et al., 2005). This suggested PAA could potentially be produced through the oxidation of PAAld. Consistent with this hypothesis, Zhang *et al*. found that *E. coli* aldehyde dehydrogenase H (AldH) can convert PAAld to PAA (Zhang et al., 2017). Given our previous biochemical studies showing that *Pto*DC3000 AldA converts IAAld to IAA (McClerklin et al., 2018), we wondered if *Pto*DC3000 can synthesize PAA and, if so, whether *Pto*DC3000 AldA (or related aldehyde dehydrogenases) might be involved.

To investigate this, we conducted feeding studies with potential PAA precursors and quantified PAA levels after 48 hrs of growth. Bacterial cultures were grown in HSC supplemented with either 100 µM phenylalanine or 25 µM PAAld. Aldehydes can be inherently toxic; we determined 25 µM PAAld to be the highest concentration of compound that did not significantly inhibit the growth of wild-type *Pto*DC3000 (WT) (Figure S1). These tests revealed significantly higher PAA in cultures fed with PAAld compared to controls (Figure 1A). In contrast, PAA levels in cultures fed with phenylalanine were slightly, but not significantly higher than in controls. The low level of PAA produced by feeding with phenylalanine suggests *Pto*DC3000 is unable to efficiently convert the amino acid to PAAld. Thus, we surmise PAA production in *Pto*DC3000 via PAAld is the predominant pathway in culture.

**Figure 1.**
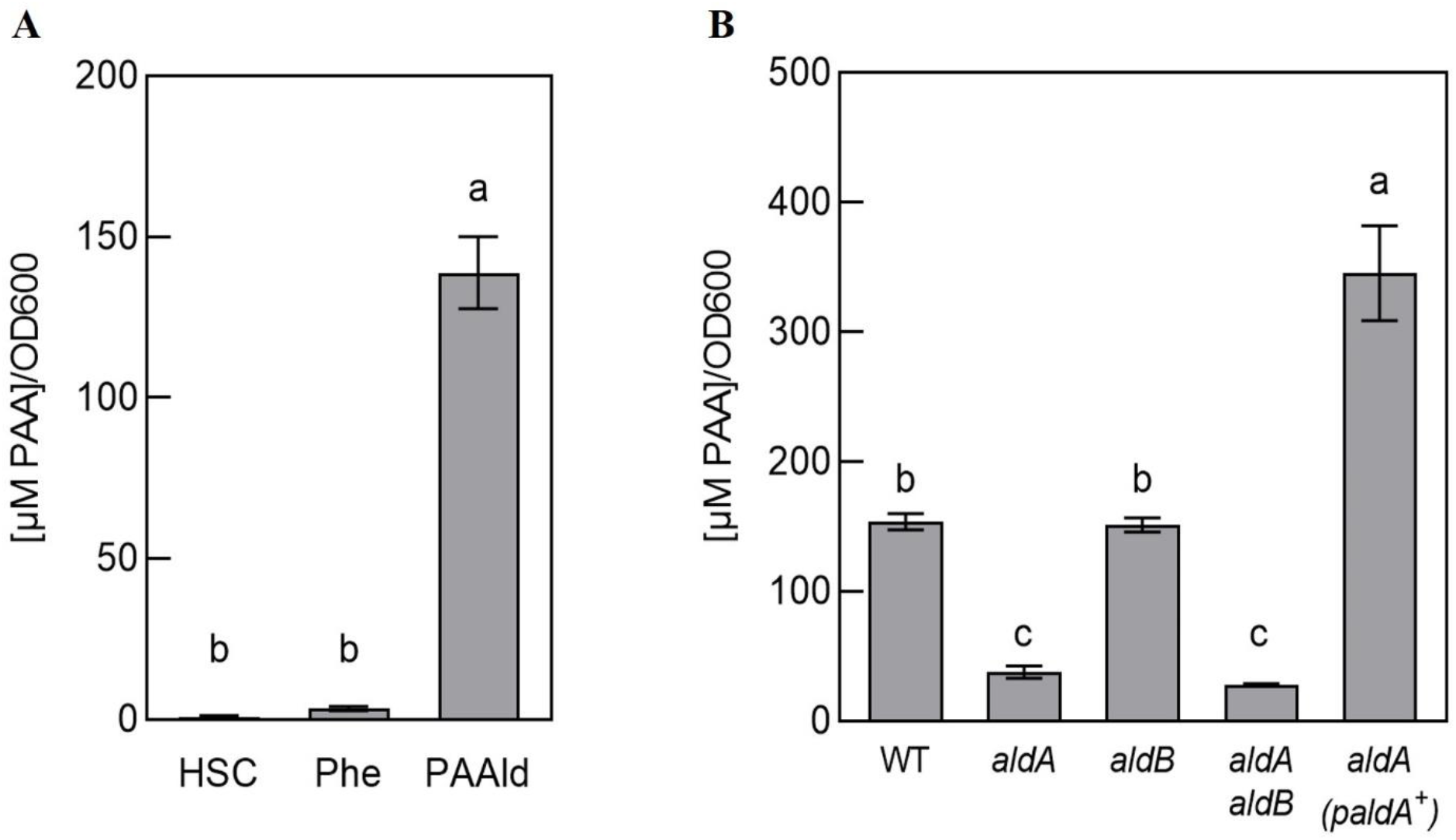
*Pto*DC3000 synthesizes phenylacetic acid (PAA) in culture. **(A)** Quantification of PAA in wild-type *Pto*DC3000 cultures, 46-48 hours(hrs) after transferring cells to HSC, HSC supplemented with 100 µM phenylalanine or 25 µM phenylacetylaldehyde (PAAld). PAA levels were measured using LC-MS/MS. Values are combined from 3 independent experiments with 3 biological replicates each (n=9) and shown as mean ± SEM. Lowercase letters indicate significant differences between treatments as determined by ANOVA followed by Tukey’s HSD test (*p* < 0.05). **(B)** Quantification of PAA in wild-type *Pto*DC3000 (WT), *aldA*::Ω, *aldB, aldA*::Ω *aldB* mutant strains or the *aldA*::Ω mutant carrying the wt *aldA* gene on a plasmid (*paldA*^+^) . PAA levels were measured using LC-MS/MS at 46-48 hrs after transferring cells to HSC supplemented with 25 µM PAAld. Values are combined from two independent experiments with three biological replicates each (n=6) and shown as mean ± SEM. Lowercase letters indicate significant differences between treatments as determined by ANOVA followed by Tukey’s HSD test (*p* < 0.05).

### 3.2 PAA synthesis is dependent upon *aldA*

Of the three identified *Pto*DC3000 aldehyde dehydrogenases capable of oxidizing IAAld to IAA (AldA, AldB, and AldC) (McClerklin et al., 2018), AldA (PSPTO_0092) shares the highest similarity to *E* . *coli* AldH (73%). To determine if PAA biosynthesis in *Pto*DC3000 is dependent on AldA, we tested if an *aldA* mutant can produce PAA when fed with PAAld. The original *aldA* mutant (*aldA*::pJP5603) characterized by McClerklin et al. (McClerklin et al., 2018) was generated by plasmid insertion and is thus potentially unstable when grown in the absence of antibiotic selection. To carry out feeding studies in the absence of antibiotics, we generated a new, more genetically stable, marker replacement mutant *aldA*::Ω (see materials and methods). We verified the *aldA*::Ω mutant to be essentially identical to the original *aldA*::pJP5603 mutant in phenotype; growth of the *aldA*::Ω mutant strain was indistinguishable from WT *Pto*DC3000 in HSC media (Figure S1). The ability of this mutant to produce IAA in culture supplemented with IAAld was monitored as described previously (McClerklin et al., 2018). Consistent with the description of the original *aldA*::pJP5603 mutant (McClerklin et al., 2018), IAA was significantly reduced in the *aldA*::Ω mutant compared to wild-type *Pto*DC3000 (WT) (Figure S2). From here on, we refer to the *aldA*::Ω mutant simply as *aldA*.

To determine if AldA contributes to *Pto*DC3000 PAA synthesis in culture, we monitored the ability of the *aldA* mutant to produce PAA when fed with PAAld. We observed a 70-80% reduction in PAA levels in the mutant compared to WT (Figure 1B). The reduced PAA synthesis phenotype was complemented by introducing the wild-type *aldA* gene on a plasmid (Figure 1B). To determine whether the related AldB enzyme also contributes to PAA production, we used the *aldB* mutant (*aldB*) described previously (McClerklin et al., 2018) and a newly generated *aldA aldB* double mutant (see materials and methods). The levels of PAA produced by *aldB* were not significantly different from WT. Further, PAA synthesis by the *aldA aldB* double mutant was not significantly lower than the *aldA* single mutant (Figure 1B). These results suggest that *aldB* is not involved in producing PAA. Thus, AldA is responsible for the majority of PAA produced by *Pto*DC3000 in culture when fed with PAAld. It is unclear where the small amount of PAA (∼ 25% WT levels) that accumulated in the supernatant of the *aldA* and *aldA aldB* mutants comes from.

### 3.3 PAAld is a substrate for AldA

Aldehyde dehydrogenases from *Pto*DC3000 have previously been biochemically and genetically examined to determine their roles in both IAA biosynthesis and in virulence (Lee et al., 2020; McClerklin et al., 2018; Zhang et al., 2020). To further investigate the biochemical basis for how AldA contributes to PAA biosynthesis, the substrate preference of AldA was investigated using purified AldA as previously described (McClerklin et al., 2018; Zhang et al., 2020). PAAld and a series of aromatic aldehydes, such as IAAld, hydrocinnamaldehyde, and cinnamaldehyde, were tested as substrates in enzymatic assays (Figure 2A). In substrate screening experiments with saturation concentrations of substrate (1 mM) and cofactor NAD^+^ (5 mM); IAAld, the known substrate of AldA (McClerklin et al., 2018), resulted in the highest activity (Figure 2B). PAAld showed the next highest activity, reaching ∼90% of the activity of IAAld. AldA had relatively low activity (50%) for hydrocinnamaldehyde and no activity for cinnamaldehyde. Steady-state kinetic analysis of AldA with PAAld and NAD^+^ confirmed that PAAld is another highly preferred substrate for AldA (Table 1). In the presence of variable concentrations of PAAld, AldA followed Michaelis-Menten kinetics, with the turnover rate (*k*_cat_) of 1124 ± 78 min^-1^ for PAAld and 83.8 ± 1.1 min^-1^ for NAD^+^, respectively. Compared to previously reported kinetic parameters for IAAld, the *k*_cat_ of PAAld was about five-fold higher, and the catalytic efficiency (*k*_cat_/*K*_m_) of PAAld was comparable at 89% (McClerklin et al., 2018). Overall, these *in vitro* substrate screening and steady-state kinetic analyses suggest that AldA can accept a range of aromatic aldehyde substrates but shows a distinct preference for PAAld.

**Table 1.** Steady-state kinetic analysis of AldA with PAAld.

| **Protein** | **Substrate** | **$k_{\text{cat}}$ (min<sup>-1</sup>)** | **$K_{\text{m}}$ (mM)** | **$k_{\text{cat}}/K_{\text{m}}$ (M<sup>-1</sup> s<sup>-1</sup>)** |
| --- | --- | --- | --- | --- |
| AldA | PAald | 1124 ± 78 | 645 ± 113 | 29,040 |
| AldA | NAD<sup>+</sup> | 83.8 ± 1.1 | 11.2 ± 1.2 | 125,020 |

Assays were performed as described in the Methods.
Average values are expressed as a mean ± SEM (n=3).

**Figure 2.**
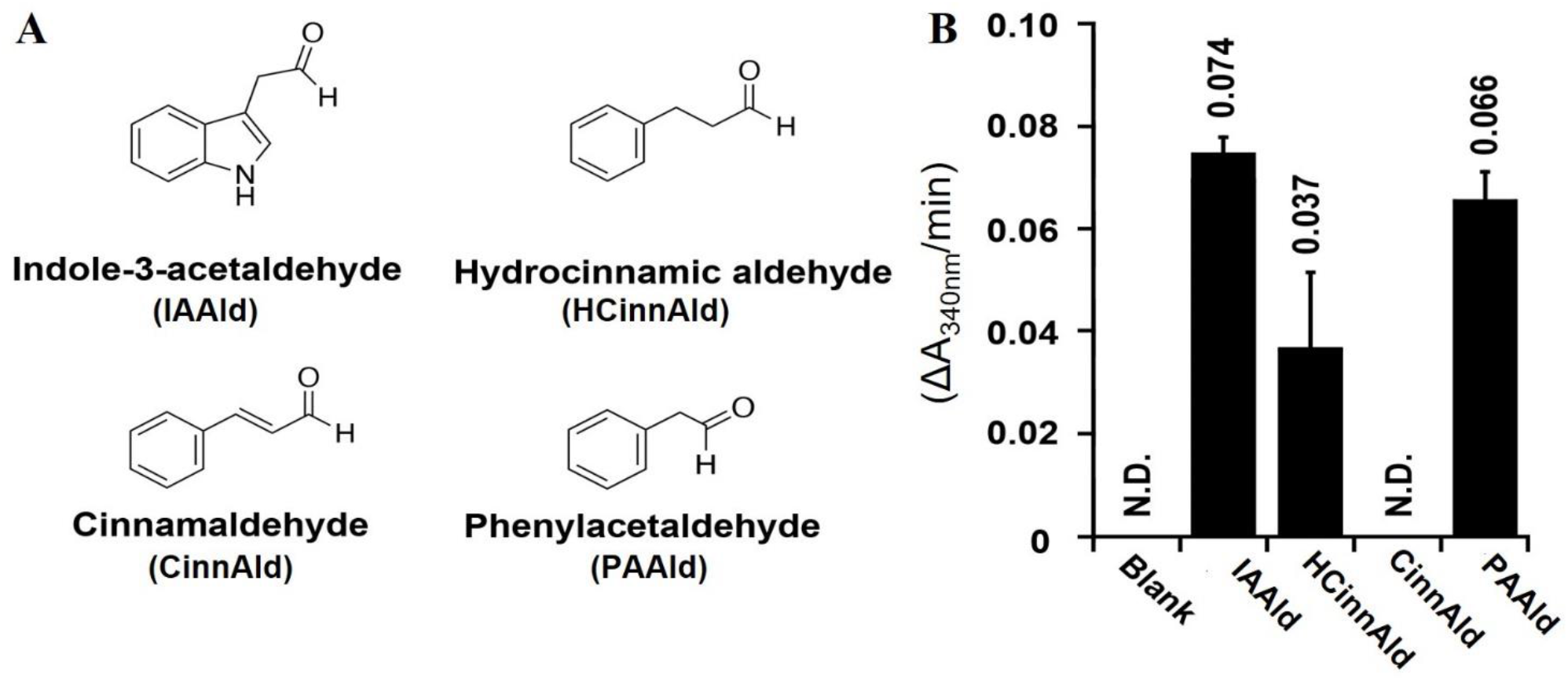
AldA can use several phenolic aldehyde substrates in vitro. **(A)** Chemical structures of aromatic aldehydes indole-3-acetaldehyde (IAAld), hydrocinnamaldehyde (HCinnAld), cinnamaldehyde (CinnAld), and phenylacetylaldehyde (PAAld) used in substrate screening. **(B)** AldA activity with the indicated aromatic aldehyde substrates. Assays were performed as described in Methods. Enzymatic activity was measured spectrophotometerically (A_340nm_) with 1 mM of NAD^+^ and 5 mM of the indicated aldehyde. Spectrophotometric absorbance changes versus time (ΔA_340nm_/min) are plotted as bar graphs for AldA-catalyzed conversion of IAAld, HCinnAld, CinnAld, and PAAld.

To provide insight into understanding how AldA accommodates PAAld in the active site, the two most preferred substrates, PAAld and IAAld, were computationally docked into the active site of the crystal structure of AldA (PDB: 5IUW) in the presence of NAD^+^ to form a ‘dead-end’ complex (Figure S3B and S3C). Computational docking of PAAld and IAAld into the active site produced models that were structurally similar to the experimentally determined structure, and the calculated receptor-ligand binding affinities were in the same range: −5.3 kcal mol^-1^ for PAAld and −5.8 kcal mol^-1^ for IAAld (Figure S3A-C). In the PAAld docking model with the highest affinity, PAAld occupies a position similar to that of IAAld with the indole moiety of IAAld replaced by its phenyl moiety. Similar to the way IAAld is positioned in the hydrophobic substrate-binding pocket, the phenyl moiety of PAAld forms multiple aromatic (*i*.*e*., Phe169, Trp176, Phe296, Trp454, and Phe467) and nonpolar interactions (*i*.*e*., Val119, Met172, Met173, and Val301) with the amino acid cluster of the AldA active site, including a π-stacking interaction with Phe169 (Figure S3D) (Lee et al., 2020; McClerklin et al., 2018). The reactive aldehyde group of each substrate is located near Cys302, the conserved catalytic cysteine (Figure S3B-D). This suggests that the catalytic activity of AldA with PAAld is due to the nonpolar surface-ligand interaction, as well as the proper accessibility of the catalytic cysteine to the reactive aldehyde group of PAAld.

### 3.4 AldA-dependent auxin synthesis in *Pto*DC3000 does not contribute to elevated auxin levels in infected plant tissue

Previous studies reported that IAA levels increase in *Pto*DC3000-infected *A. thaliana* leaves (Chen et al., 2007). Given that *Pto*DC3000 can synthesize IAA (McClerklin et al., 2018) and PAA (Figure 1) via the AldA-dependent pathway in culture, we explored the possibility that the contribution of *Pto*DC3000 AldA activity to the observed increase in auxin levels in infected *A. thaliana* leaves by quantifying IAA, PAA, and their amino acid conjugates in *Pto*DC3000-infected leaf tissues. The leaves were infiltrated with wild-type *Pto*DC3000 (WT) or the *aldA*::Ω mutant strain (*aldA*) and collected at 24 and 48 hrs post infection (hpi) for auxin quantification. Consistent with prior reports, we observed a 3.2-fold increase in IAA levels in WT-infected leaves compared to mock treatment at 48 hpi (Figure 3A) (Chen et al., 2007). Notably, *aldA*-infected leaves exhibited a comparable increase in IAA levels at 48 hpi (Figure 3A), indicating that the AldA activity of *Pto*DC3000 does not significantly contribute to IAA accumulation after infection.

**Figure 3.**
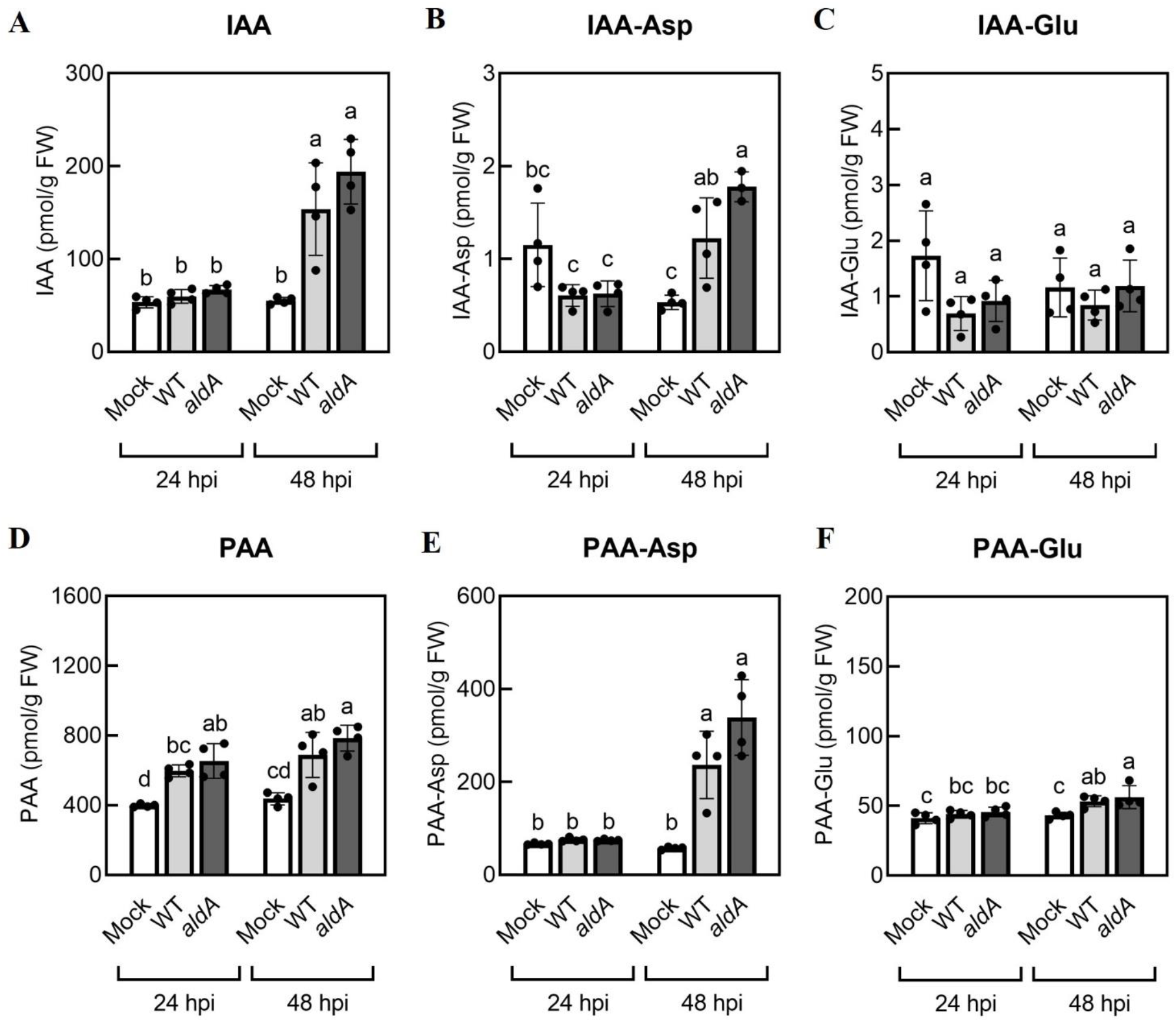
Auxin levels increase in *A. thaliana* plants inoculated with *Pto*DC3000. **(A)** Indole-3-acetic acid (IAA), **(B)** IAA-aspartate conjugate (IAA-Asp), **(C)** IAA-glutamate conjugate (IAA-Glu), **(D)** phenylacetic acid (PAA), **(E)** PAA-aspartate conjugate (PAA-Asp), **(F)** PAA-glutamate conjugate (PAA-Glu). Five-week-old wild-type *A. thaliana* plants (Col-0) were infiltrated with 10 mM MgCl_2_ (mock), wild-type *Pto*DC3000 (WT), or the *aldA*::Ω (*aldA*) mutant. The inoculum used for infiltration was ∼1 × 10^6^ CFU/mL. Infiltrated leaves were collected for LC-MS/MS analysis of auxin metabolites at 24 and 48 hrs post inoculation (hpi). Data are from one representative experiment (n=4) and shown as mean ± SD. Similar results were obtained in two additional independent experiments. Lowercase letters indicate significant differences between samples as determined by ANOVA followed by Tukey’s HSD test (*p* < 0.05). FW: fresh weight of leaf tissues.

Plants maintain tight control over free auxin levels and auxin-conjugates form rapidly in response to increases in auxin (Korasick et al., 2013). Thus, in addition to free IAA, we also monitored the levels of the IAA-amino acid conjugates IAA-aspartate (IAA-Asp) and IAA-glutamate (IAA-Glu) in the infected leaf tissue. IAA-Asp and IAA-Glu were detectable in infected leaves but were present at substantially lower levels compared to free IAA (Figure 3B and 3C). Specifically, IAA-Asp exhibited a small but significant increase (∼2.3-fold) at 48 hpi in WT-and *aldA-*infected leaves compared to mocked leaves (Figure 3B). No change in IAA-Glu was detected in infected tissue. As with free IAA, leaves inoculated with the *aldA* mutant did not accumulate significantly different levels of IAA-amino acid conjugates compared to leaves infected with WT. Thus, it appears that IAA synthesis by *Pto*DC3000 does not significantly contribute to the increase in IAA levels in infected plant tissue. Alternatively, if *Pto*DC3000 IAA synthesis plays a role in IAA accumulation, it is via a mechanism independent of AldA activity.

Consistent with the previous report (Sugawara et al., 2015), mock-inoculated *A. thaliana* leaves had remarkably higher PAA levels compared to IAA (Figure 3A and 3D). In two out of three independent experiments, we observed modest yet statistically significant increases of PAA in WT-infected leaves by 24 hpi (ranging from a 1.2-to 1.9-fold increase), and in all three experiments, we consistently observed significantly elevated PAA levels by 48 hpi (ranging from a 1.5-to 1.8-fold increase, Figure 3B). There was no difference in accumulation of PAA between leaves inoculated with WT and *aldA* strains at either time point. Further quantification of PAA-amino acid conjugates showed significant increases in the levels of both PAA-Asp and PAA-Glu at 48 hpi in inoculated leaves (4.7-fold and 1.2-fold increase, respectively, Figure 3E and 3F); however, as for free PAA, there was no difference between leaves inoculated with WT and *aldA* strains. We thus conclude that AldA-dependent synthesis of PAA by *Pto*DC300 does not significantly contribute to PAA accumulation in infected plant leaves.

### 3.5 Investigating the role of PAA in pathogenesis

Plants with altered IAA levels or auxin sensitivity exhibit varying degrees of susceptibility to virulent *P. syringae*. For example, plants with elevated IAA and/or IAA-Asp or increased IAA sensitivity are more susceptible to *Pto*DC3000 (Chen et al., 2007; Djami-Tchatchou et al., 2020; González-Lamothe et al., 2012; Mutka et al., 2013); whereas plants with decreased auxin sensitivity are less prone to *P. syringae* infection (Djami-Tchatchou et al., 2020; Navarro et al., 2006; Wang et al., 2007). Given our observation of increased PAA and PAA-Asp levels in infected leaves (Figure 3), we hypothesized that PAA, like IAA, could promote the pathogenesis of *Pto*DC3000 *in planta*.

To test this hypothesis, we took advantage of previously described mutant and transgenic *A. thaliana* lines with altered PAA levels. The *AROGENATE DEHYDRATASE (ADT*) gene family encodes enzymes that catalyze the conversion of arogenate to phenylalanine and have been shown to regulate the levels of PAA in *A. thaliana* (Aoi et al., 2020b) (Figure 4A). Aoi et al. (2020a) demonstrated that transgenic plants overexpressing *ADT4* or *ADT5* (*ADT4* OE or *ADT5* OE) accumulated elevated levels of phenylalanine, PAA, and PAA-amino acid conjugates in seedlings. In contrast, the *adt1 adt3 adt4 adt5 adt6* quintuple knockout mutant (*adt1/3/4/5/6*), which carries T-DNA insertions in five of the known *ADT* genes, accumulated reduced amounts of phenylalanine, PAA and PAA-Asp (Aoi et al., 2020b).

**Figure 4.**
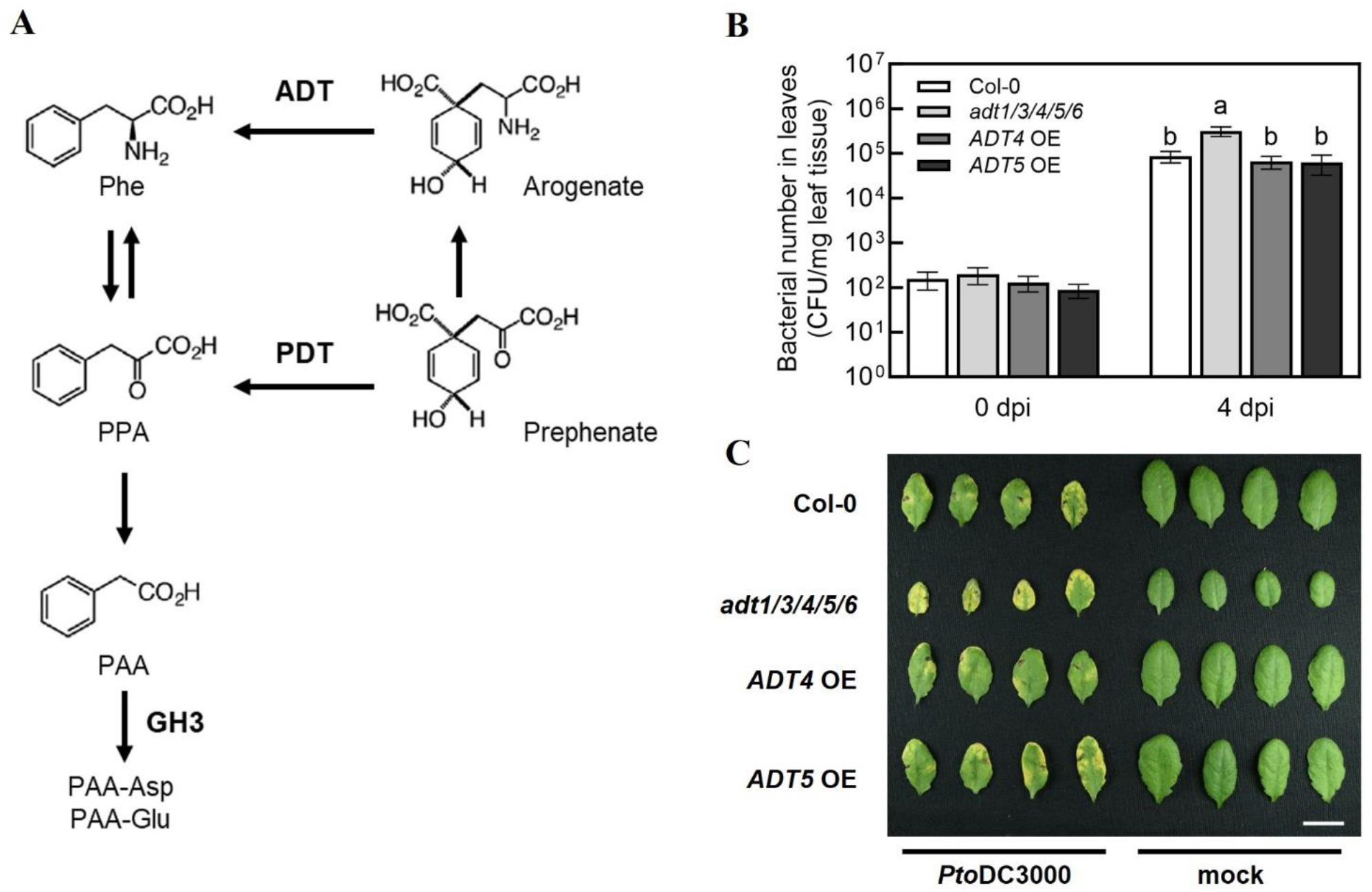
The *A. thaliana adt1/3/4/5/6* quintuple mutant exhibits increased susceptibility to *Pto*DC3000. **(A)** PAA biosynthetic and metabolic pathways in *A. thaliana*. PAA is produced from phenylalanine (Phe) via phenylpyruvate (PPA) by transamination and decarboxylation. Two dehydratases, arogenate dehydratase (ADT) and prephenate dehydratase (PDT), mediate the production of PPA, a precursor of PAA biosynthesis. The figure was modified from Aoi et al. (2020a). **(B)** Bacterial growth of wild-type *Pto*DC3000 in *A. thaliana adt* mutants and transgenic plants overexpressing *ADT4* or *ADT5*. Five-week-old wild-type *A. thaliana* (Col-0), the *adt1 adt3 adt4 adt5 adt6* quintuple mutant (*adt1/3/4/5/6*), *ADT4* overexpressing (*ADT4* OE) and *ADT5* overexpressing (*ADT5* OE) plants were infiltrated with ∼1 × 10^5^ CFU/mL of wild-type *Pto*DC3000. Bacterial growth in infiltrated leaves was quantified 0-and 4-day post-inoculation (dpi). Data are combined from three independent experiments and shown as mean ± SD (n=12 for 0 dpi, n=24 for 4 dpi). Letters indicate significant differences between genotypes on day 4 as determined by ANOVA followed by Tukey’s HSD test (*p* < 0.05). **(C)** Disease symptoms of *A. thaliana* leaves 4 dpi. Plants of the indicated genotypes were infiltrated with ∼1 × 10^6^ CFU/mL of wild-type *Pto*DC3000. Leaves infiltrated with 10 mM MgCl_2_ (mock) are shown on the right. The photograph is from one representative experiment. Scale bar indicates 1 cm. CFU: Colony forming units; FW: fresh weight of leaf tissue.

Based on our initial hypothesis, we anticipated that *adt1/3/4/5/6* mutant plants would exhibit reduced susceptibility to *Pto*DC3000, while *ADT4* OE and *ADT5* OE lines would be more susceptible. We infiltrated WT *Pto*DC3000 into five-week-old wild-type *A. thaliana* (Col-0), *adt1/3/4/5/*6, *ADT4* OE, and *ADT5* OE plants, and quantified bacterial growth at 0-and 4-day post inoculation (dpi). As expected, WT grew three orders of magnitude in Col-0 (Figure 4B). Surprisingly, *adt1/3/4/5/6* plants supported slightly higher levels of bacterial growth and exhibited more severe disease symptoms compared to Col-0 (Figure 4B and 4C). On the other hand, *ADT4* OE and *ADT5* OE plants did not exhibit any alteration in susceptibility compared to Col-0. Thus, contradictory to our initial predictions, plants with reportedly reduced PAA levels are more susceptible to *Pto*DC3000, and plants previously shown to have elevated PAA levels exhibited WT susceptibility.

One plausible explanation for these results could be that the PAA levels in mature plants grown under our conditions were not altered as described in the literature. In our plant infections, we used five-week-old plants grown on soil with a short-day photoperiod at 21°C. Conversely, plants used for published auxin quantification were ten-day-old seedlings grown on sterile Murashige and Skoog (MS) agar under a long-day photoperiod at 23°C (Aoi et al., 2020b). Given the potential influence of both developmental stage and environment on phytohormone levels, we wondered if these mutant plants accumulated altered levels of PAA and PAA-amino acid conjugates under our experimental conditions.

To investigate this, we quantified free PAA and PAA-amino acid conjugates in uninfected five-week-old *adt1/3/4/5/*6 mutant and *ADT* OE transgenic lines grown under our experimental conditions. The *adt1/3/4/5/*6 plants accumulated essentially wild-type levels of PAA (Figure 5A) but significantly reduced levels of PAA-Asp and PAA-Glu (Figure 5B and 5C). In contrast, the *ADT4 OE* and *ADT5 OE* plants accumulated wild-type levels of all PAA forms (Figure 5A-C), rather than the anticipated elevated levels. These data suggest that PAA levels are dependent on developmental stage and/or growth conditions. This could potentially explain why we did not see altered susceptibility to *Pto*DC3000 in *ADT* overexpression lines. Furthermore, these data introduce an alternative hypothesis regarding the role of PAA in plant susceptibility. That is, as opposed to free PAA promoting pathogenesis, PAA-Asp and PAA-Glu may negatively regulate plant susceptibility to *Pto*DC3000.

**Figure 5.**
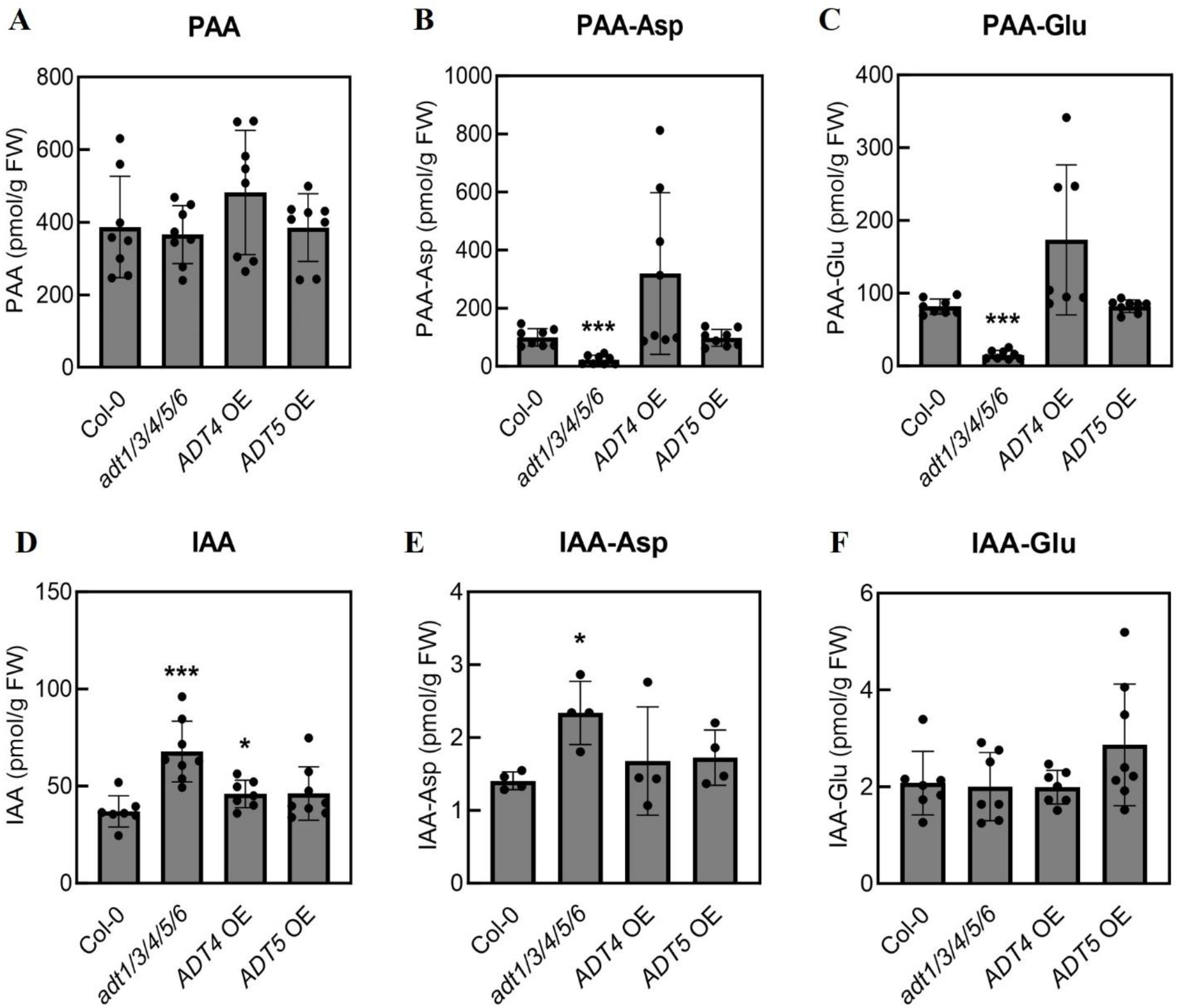
Mature *adt1/3/4/5/6* mutant plants accumulate reduced levels of PAA-amino acid conjugates but increased levels of IAA and IAA-Asp. Quantification of **(A)** phenylacetic acid (PAA), **(B)** PAA-aspartate conjugate (PAA-Asp), **(C)** PAA-glutamate conjugate (PAA-Glu), **(D)** indole-3-acetic acid (IAA), **(E)** IAA-aspartate conjugate (IAA-Asp), and **(F)** IAA-glutamate conjugate (IAA-Glu) in uninoculated *A. thaliana* plants. Leaves from five-week-old wild-type *A. thaliana* (Col-0), *adt1/3/4/5/6* quintuple mutant, *ADT4* overexpressing (*ADT4* OE) and *ADT5* overexpressing (*ADT5* OE) plants were collected for LC-MS/MS analysis of PAA metabolites. Data are combined from two independent experiments with four biological replicates each (n=8) and shown as mean ± SD. Asterisks indicate significant differences between mutant or transgenic lines and Col-0 as determined by Student’s *t*-test (^*^: *p* < 0.05; ^* *^: *p* < 0.01; ^* * *^: *p* < 0.001). FW: fresh weight of leaf tissues.

Previous studies on the interplay of the different forms of auxins in *A. thaliana* have revealed that PAA and PAA-amino acid conjugates can impact IAA homeostasis (Perez et al., 2023). Thus, we hypothesized that the increased susceptibility of *adt1/3/4/5/*6 might be attributable to altered IAA levels in the plants. We quantified IAA and its conjugates in these lines, and indeed, the levels of IAA and its conjugates were altered in *adt1/3/4/5/*6. The *adt1/3/4/5/*6 plants accumulated significantly higher levels of IAA and IAA-Asp compared to Col-0 (Figure 5D and 5E), which are correlated with increased susceptibility to *Pto*DC3000 (Figure 4B). Notably, the *adt1/3/4/5/6* mutant plants were much smaller than wild-type plants (Figure 4C), but did not show any other phenotypes typical for plants with elevated IAA levels. Thus, the reduced growth phenotypes of the mutant could be due to reduced phenylalanine levels (Aoi et al., 2020b). We did not observe any significant increase in PAA or PAA-amino acid conjugates, nor changes in free or amino acid-conjugated IAA levels in the *ADT OE* lines (Figure 5). This is consistent with our findings that the *ADT OE* plants did not exhibit altered susceptibility to *Pto*DC3000 (Figure 4). In summary, our results suggest that PAA-amino acid conjugates and/or the regulatory crosstalk between these conjugates and IAA mediate *A. thaliana* susceptibility to *Pto*DC3000.

## 4 Discussion

To further our understanding of the roles of auxins during pathogenesis of *P. syringae*, we investigated whether *Pto*DC3000 can synthesize PAA and if this synthesis influences the outcome of infection of *A. thaliana*. We also took advantage of an *A. thaliana* mutant line compromised for PAA synthesis to assess the impact of reduced endogenous levels of PAA-related molecules on disease susceptibility.

### 4.1 *Pto*DC3000 synthesizes PAA in culture, using PAAld as a substrate

We demonstrated that *Pto*DC3000 can synthesize PAA in culture when fed with PAAld, and that PAA production largely depends on the aldehyde dehydrogenase AldA (Figure 1). In these experiments, PAA accumulation in the *aldA* mutant culture was reduced to about 25% of WT levels, and introduction of the *aldB* mutation did not further reduce PAA production. This suggests that *Pto*DC3000 may encode one or more additional enzymes with PAAld dehydrogenase activity. Alternatively, the residual amount of PAA present in these cultures could be due to conversion of PAAld to PAA via an activity that accumulates in the medium during the growth of the bacterium. To investigate this, we incubated PAAld in conditioned HSC medium for 48 hrs and then quantified PAA levels. The molar concentration of PAA measured in conditioned HSC made from either WT or the *aldA* mutant after incubation with PAAld (∼21-32 µM) was similar to the starting concentration of PAAld (25 µM), whereas only very small amounts of PAA were detected in non-conditioned HSC that had been incubated with PAAld (Table S4). These observations support the hypothesis that an AldA-independent activity that accumulates in the media after prolonged growth of *Pto*DC3000, can convert PAAld into PAA.

The observation that *Pto*DC3000 did not synthesize PAA when fed with phenylalanine (Figure 1A) suggests that *Pto*DC3000 is unable to convert the amino acid to PAAld (*i*.*e*., via the intermediate phenylpyruvate, PPA). This is consistent with the observation that the *Pto*DC3000 genome does not encode an obvious phenylpyruvate decarboxylase, an enzyme found in other bacteria that catalyzes the decarboxylation of PPA to PAAld (Patten et al., 2013; Spaepen et al., 2007). Although we cannot formally rule out that the inability to use phenylalanine as a substrate for PAA is due to the inability to take up the amino acid from growth media, we think this is unlikely, as the *Pto*DC3000 genome is predicted to encode at least one aromatic amino acid transporter homologous to the AroP1 transporter of *P. aeruginosa* (www.pseudomonas.com).

### 4.2 AldA is a versatile enzyme that can use structurally similar substrates

AldA was initially identified as an enzyme that catalyzes the conversion of IAAld to IAA in *Pto*DC3000 (McClerklin, 2018). Here, we show that this enzyme uses a broad spectrum of aromatic substrates. Specifically, it accepts both IAAld and PAAld to synthesize two distinct auxin species, IAA and PAA, respectively (Figure 2B). The substrate promiscuity of enzymes in auxin biosynthesis has been documented in studies on higher plants. For instance, to examine the conversion of IAAld to IAA by aldehyde oxidase (AO; EC 1.2.3.1), three aldehyde oxidase homologs (*i*.*e*., AO1, AO2, and AO3) from *A. thaliana* were tested against a selection of 11 distinct aldehydes, including IAAld and PAAld (Seo et al., 1998). While AO1 exhibited a strong substrate preference for IAAld and indole-3-aldehyde (IAld), all three AOs showed activity across a broad spectrum of aromatic aldehydes. Despite the lack of amino acid sequence, structural, and/or mechanistic similarity between the plant AOs and *Pto*DC3000 AldA, both types of enzymes exhibit the capacity to accommodate a wide variety of aromatic compounds.

Previous revelation of the three-dimensional structure of the AldA•NAD^+^•IAA complex indicates that the IAAld/IAA binding site of AldA is predominantly formed by amino acid residues with high hydrophobicity, suggesting apolar interactions are the primary binding mechanism (McClerklin, 2018). Particularly, the sidechains of Phe169, Phe296, and Phe467 may be involved in potential π-π interactions with the indole moiety of IAAld during binding. Given the substrate binding site environment of AldA and the structural similarity of the aromatic substrates, the substitution of the indole and phenyl moieties seem to minimally impact the substrate accessibility to the active site or its binding affinity for PAAld. AldA can also use another structurally similar aromatic aldehyde, hydrocinnamaldehyde, as a substrate, presumably to produce hydrocinnamic acid (a.k.a., phenylpropanoic acid). The fact that cinnamaldehyde, in which the aldehyde sidechain contains a carbon-carbon double bond, is not a good substrate for AldA may provide some clues about the substrate preference for this enzyme. As discussed below, the ability of *Pto*DC3000 to synthesize the auxins IAA and PAA is biologically relevant, and both IAAld and PAAld are readily found in plants (Gutensohn et al., 2011; Koshiba et al., 1996). However, the biological relevance of hydrocinnamaldehyde production by *Pto*DC3000 is not clear.

### 4.3 AldA-dependent auxin synthesis may participate in aspects of *Pto*DC3000 biology other than contributing to increasing auxin levels in infected plant tissues

Our finding that neither the *aldA*::Ω mutant featured in this study, nor the *aldA*::pJP5603 insertion mutant originally characterized by McClerklin *et al*. (2018) exhibited reduced growth on *A. thaliana* plants (Figure S4) suggests that AldA does not play a major role during pathogenesis on *A. thaliana*. This is consistent with our observation that AldA-dependent auxin synthesis does not appear to contribute significantly to the increase in either IAA or PAA levels in infected leaves (Figure 3). Thus, the increase in auxin appears to be due to synthesis by the plant, in response to infection; however, because endogenous auxin levels are elevated in infected leaves (3-fold and 1.5-fold increase in IAA and PAA, respectively compared to uninoculated controls), any further increase contributed by bacterial AldA-dependent synthesis may not be readily detectable. It is also possible that *Pto*DC3000 contributes to the increase in auxin via an AldA-independent process, involving a different acetaldehyde dehydrogenase or via a different biosynthetic pathway.

Given that the *aldA* gene is highly conserved in *P. syringae* strains, AldA-dependent production of PAA, IAA, or both aromatic acids may play important roles at different stages of the bacterial life cycle, such as during epiphytic growth or during communication with other microorganisms. Alternatively, AldA-dependent auxin synthesis may be involved in regulation of bacterial gene expression. For example, IAA impacts the expression of virulence-related genes in *P. syringae* (Djami-Tchatchou et al., 2022; Djami-Tchatchou et al., 2020), *Acinetobacter baumannii* (Hooppaw Anna et al., 2022), *Agrobacterium tumefaciens* (Liu and Nester, 2006), and *Erwinia chrysanthemi* (Yang et al., 2007).

### 4.4 PAA/IAA homeostasis impacts susceptibility to *Pto*DC3000

Auxin promotes disease susceptibility to many biotrophic pathogens (Kunkel and Harper, 2018). For example, exogenous application of IAA or IAA-Asp promotes disease development in plants infected with *Pto*DC3000 (Chen et al., 2007; González-Lamothe et al., 2012), and plants with elevated levels of IAA exhibit increased susceptibility (Chen et al., 2007; Cui et al., 2013; Djami-Tchatchou et al., 2020; Navarro et al., 2006). Genetic and molecular studies further reveal that IAA can promote pathogenesis via two separate mechanisms: (1) repressing salicylic acid (SA)-mediated host defense responses (McClerklin et al., 2018; Wang et al., 2007) and (2) regulating bacterial virulence gene expression (Djami-Tchatchou et al., 2020).

PAA has been shown to play similar regulatory roles as IAA in plants, and responses to PAA are mediated via the same phytohormone response system (Shimizu-Mitao and Kakimoto, 2014). We hypothesized that PAA would also serve a similar role as IAA during *Pto*DC3000 infection and promote pathogenesis. Thus, we predicted that plants reported to have elevated PAA (or PAA-amino acid conjugates) would show increased susceptibility, while plants with reduced PAA (or PAA-amino acid conjugates) would exhibit decreased susceptibility. Surprisingly, we observed that the *adt1/3/4/5/6* mutant, which accumulated reduced levels of PAA-AAs (Figure 5B and 5C), exhibited increased susceptibility (Figure 4B and 4C).

In addition to reducing PAA-Asp and PAA-Glu levels, the disruption of *ADT1/3/4/5/6* resulted in increased levels of IAA and IAA-Asp (Figure 5D and 5E). We hypothesize that the genetic modulation of the PAA pool in plants impacts IAA homeostasis, which is then responsible for the increased susceptibility. Aligning with this idea, several mechanisms have been proposed for modulating the metabolic crosstalk between IAA and PAA. These mechanisms include: (i) modulation of GH3 (GRETCHEN HAGEN 3) acyl acid amido synthetases, which can catalyze formation of aspartate and glutamate conjugated forms of IAA and PAA (Westfall et al., 2016) and/or UDP-dependent glycosyltransferase (UGT)-dependent modification of auxins (Aoi et al., 2020a; Aoi et al., 2020c) and (ii) repression of IAA biosynthesis genes by PAA (Perez et al., 2021). The first homeostasis crosstalk mechanism suggests that free PAA and/or IAA induces the activity of auxin modification enzymes, leading to the accumulation of conjugated PAA and/or IAA. This mechanism does not adequately explain our findings, given that we observe the opposite outcome in the *adt1/3/4/5/6* mutant. On the other hand, the relationship between PAA and IAA levels we observed in the *adt1/3/4/5/6* mutant could be achieved through the negative crosstalk between PAA and IAA biosynthesis. In this scenario, plants with a reduced PAA pool could result in increased synthesis of IAA. This balancing between phenylalanine-and tryptophan-derived auxins could result from the same mechanisms that regulate distribution of chorismate into either aromatic amino acid pool (Kroll et al., 2017), which may impact subsequent synthesis of PAA and IAA. We cannot rule out the possibility of direct inhibition of pathogen virulence by PAA-amino acid conjugates. For example, reduced levels of PAA-Asp and PAA-Glu could potentially promote *Pto*DC3000 virulence through IAA-independent mechanisms, such as impacting bacterial virulence gene expression or plant defense responses. Future studies investigating the roles of PAA and PAA-amino acid conjugants in modulating host defenses and/or bacterial gene expression should provide new insights into the roles of auxin in plant-microbe interactions.

## 5 Data availability statement

Not applicable.

## 6 Author contributions

CYL: Conceptualization, Experimentation, Data Analysis, Writing – original draft, review and editing; CPH: Conceptualization, Experimentation, Data Analysis, Writing – original draft, review and editing; SGL: Conceptualization, Experimentation, Data Analysis, Writing – review and editing; YQ: Experimentation, Writing – review and editing; TC: Experimentation; YA: Experimentation; JMJ: Conceptualization, Writing - review and editing; HK: Conceptualization, Experimentation, Writing – review and editing; JAVB: Conceptualization, Writing – review and editing; BNK: Project administration, Conceptualization, Writing – original draft, review and editing.

## 8 Acknowledgements

We thank Sarah A. Pardi for help characterizing the *aldA*::Ω mutant.

## 10 Supplementary material

**Figure S1:** Growth of wild-type *Pto*DC3000 and the indicated *ald* mutants in Hoitkin-Sinden medium containing 10 mM citrate (HSC) and HSC supplemented with 25 µM phenylacetylaldehyde (PAAld).

**Figure S2**. Quantification of indole-3-acetic acid (IAA) produced in culture by wild-type *Pto*DC3000 (WT), the *aldA*::Ω mutant (*aldA*), and the complemented *aldA*::Ω mutant (*aldA (pAldA*^*+*^)).

**Figure S3**. Comparison of AldA-ligand binding in the active site tunnel.

**Figure S4**. Bacterial growth of wild-type *Pto*DC3000 strains in *A. thaliana* leaves sampled for auxin quantification.

**Table S1**. Bacterial strains and plasmids used in this study.

**Table S2**. Primers used in this study.

**Table S3**. Metabolites detected by LC-MS/MS.

**Table S4**. Accumulation of phenylacetic acid (PAA) in conditioned medium in three independent experiments.

