## Supplemental Files for "Investigating the biosynthesis and roles of the auxin phenylacetic acid during *Pseudomonas syringae-Arabidopsis thaliana* pathogenesis"

**Supplemental Figure Legends**

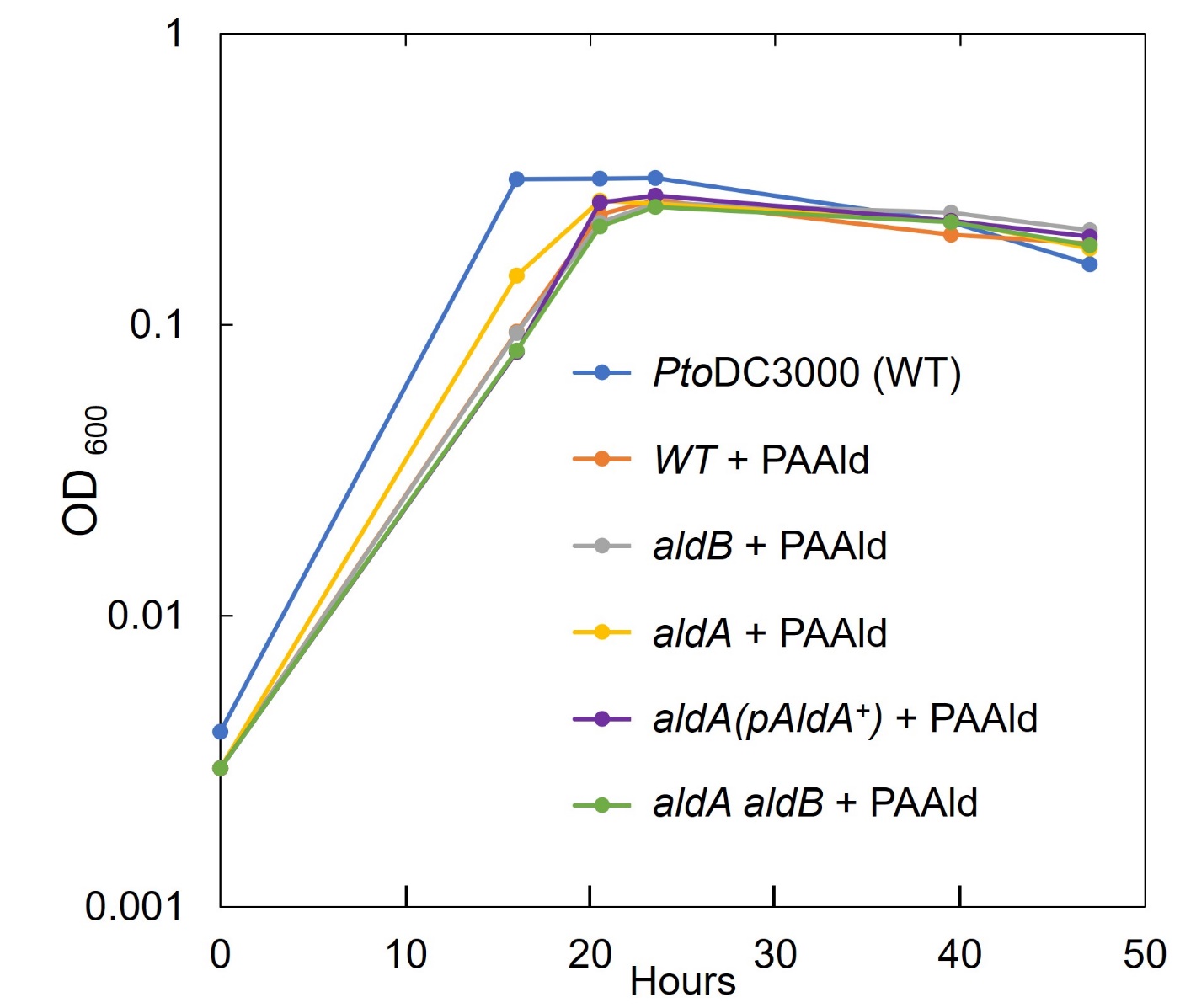

**Figure S1:** Growth of wild-type *Pto*DC3000 and the indicated *ald* mutants in Hoitkin-Sinden medium containing 10 mM citrate (HSC) and HSC supplemented with 25 µM phenylacetylaldehyde (PAAld). The data shown are from one of the experiments used to collect the data shown in Figure 1B. The strains analyzed were: wild-type *Pto*DC3000, *aldB*::pJP5603, *aldA*::Ω, *aldA*::Ω *aldB*::pJP5603 double mutant, and the *aldA*::Ω mutant carrying the pAldA^+^ complementing plasmid.

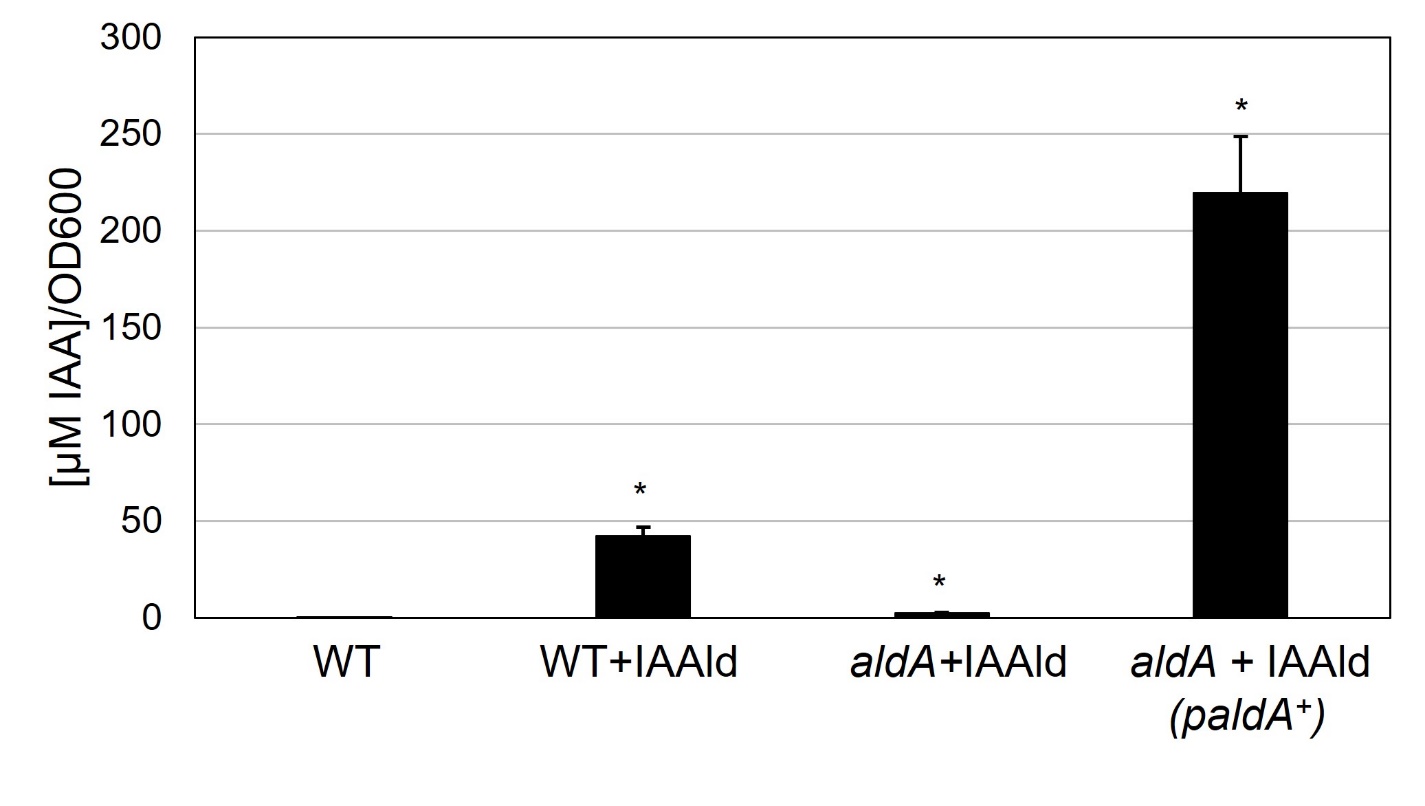

**Figure S2.** Quantification of indole-3-acetic acid (IAA) produced in culture by wild-type *Pto*DC3000 (WT), the *aldA*::Ω mutant (*aldA*), and the complemented *aldA*::Ω mutant (*aldA (pAldA^+^*)). IAA levels were measured using LC-MS/MS 46-48 hours after growing in HSC supplemented with 250 µM IAAld. Data are from 3 biological replicates (n=3) and shown as mean ± SEM. Asterisks indicate significant differences of IAA accumulation in cultures between WT without IAAld supplementation and indicated strains supplemented with IAAld as determined by Student’s *t*-test (*: *p* < 0.05).

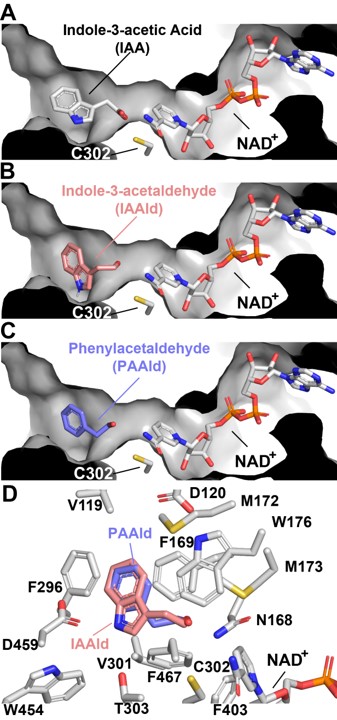

**Figure S3.** Comparison of AldA-ligand binding in the active site tunnel. **(A)** Indole-3-acetic acid (IAA) (product, PDB entry ID: 5IUW), **(B)** indole-3-acetaldehyde (IAAld) (substrate, in rose), **(C)** phenylacetylaldehyde (PAAld) (substrate, in blue), and NAD^+^ (cofactor) are positioned through molecular docking experiments and shown as stick models. The active site tunnel is shown as a surface view. **(D)** Substrate binding site. Side-chains of residues interacting with IAAld (in rose) and PAAld (in blue) are shown as stick-renderings.

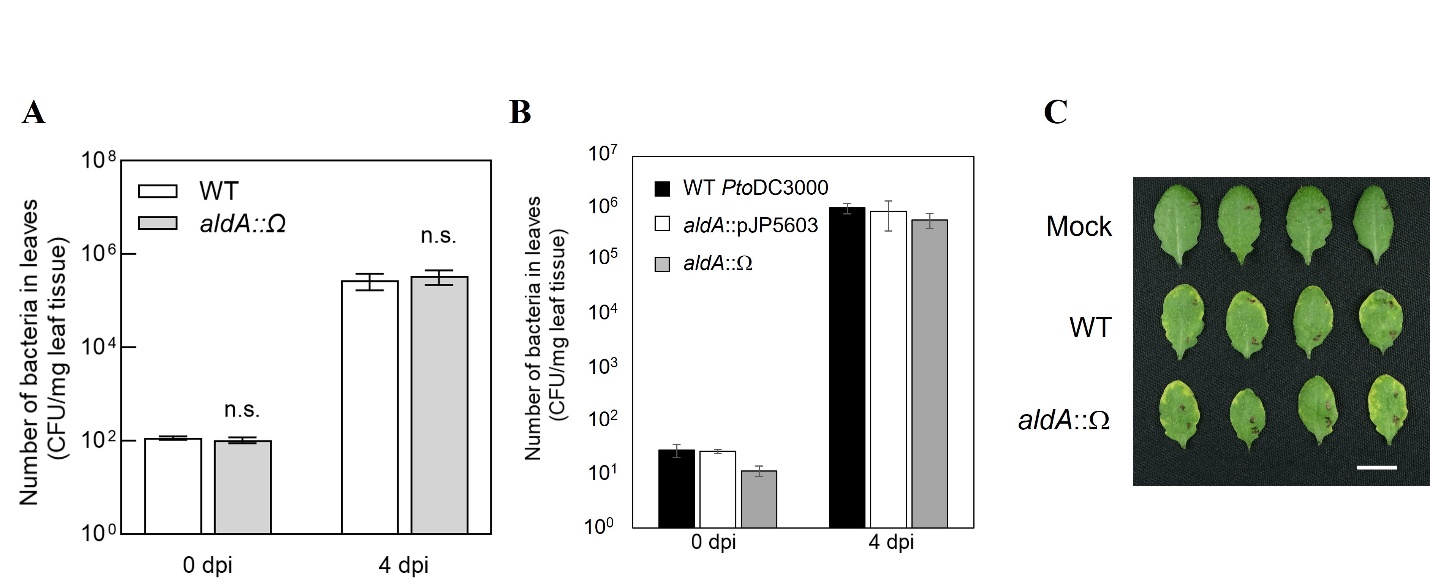

**Figure S4.** Bacterial growth of wild-type *Pto*DC3000 strains in *A. thaliana* leaves sampled for auxin quantification. **(A)** Bacterial growth of wild-type *Pto*DC3000 (WT), and *aldA* insertion mutant (*aldA*::Ω) in inoculated leaves was quantified at ~4 hours after inoculation (day 0) and 4 days post-inoculation (dpi). Data from one representative experiment are shown as mean ± SD (n=4 for 0 dpi and n=8 for 4 dpi). Similar results were obtained in two independent experiments. No statistical significance (n.s.) between *Pto*DC3000 strains using Student’s *t*-test (*p* < 0.05). **(B)** Growth of WT and two independent *aldA* mutants (*aldA*::pJP5603, McClerklin et al., 2018) and *aldA*::Ω ) 0 and 4 dpi. Data from one representative experiment are shown as mean ± SD (n=4 for 0 dpi and n=8 for 4 dpi). No statistical significance between *Pto*DC3000 strains using Student’s *t*-test (*p* < 0.05). FW: fresh weight of leaf tissues. **(C)** Disease symptoms on Col-0 leaves infiltrated with 10 mM MgCl_2_ (Mock) and 10^6^ CFU/mL of indicated *Pto*DC3000 strains were photographed on 3 dpi. The photograph is from one representative experiment. Scale bar indicates 1 cm. CFU: Colony forming units; FW: fresh weight of leaf tissue.

**Supplemental Tables**

**Table S1.** Bacterial strains and plasmids used in this study.

| **Strain/Plasmid** | **Characteristics** | **Reference or Source** |
| --- | --- | --- |
| ***P. syringae* strains** | | |
| *Pto*DC3000 (WT) | Derivative of NCPPB1106; Rif^r^ | (Cuppels Diane, 1986) |
| *aldA*::Ω (*aldA*) | *PSPTO_0092* gene replaced with the omega fragment from pHP45; Rif^r^, Spec^r^ | This study |
| *aldB*::pJP5603-Km | *PSPTO_2673* disrupted by pJP5603; Rif^r^, Km^r^ | (McClerklin et al., 2018) |
| *aldA*::Ω *aldB*::pJP5603-Km | *aldA aldB* double mutant; Rif^r^ Spec^r^, Km^r^ | This study |
| *aldA*::Ω (pAldA^+^) | *aldA* mutant with complementing clone | This study |
| *aldA*::pJP5603-Ω*aldA* | Merodiploid carrying pJP5603-Ω*aldA* plasmid at *PSPTO_0092*; generated by single cross over; Intermediate strain used to create *aldA*::Ω; Rif^r^, Spec^r^, Km^r^ | This study |
| *aldA*::pJP5603-Km | *PSPTO_0092* disrupted with pJP5603; Rif^r^, Km^r^ | (McClerklin et al., 2018) |
| *aldB*::pJP5603-Tet | *PSPTO_2673* disrupted with pJP5603-Tet; Rif^r^, Tet^r^ | (McClerklin et al., 2018) |
| *E. coli* strains | | |
| DH5α λpir | *recA*, *lacZΔM15*, *λpir* | (Miller and Mekalanos, 1988) |
| MM294A | Triparental mating helper strain | (Finan et al., 1986) |
| Plasmids | | |
| pRK2013 | Helper plasmid; Cm^r^ | (Finan et al., 1986) |
| pJP5603-Km | Suicide vector; Km^r^ | (Penfold and Pemberton, 1992) |
| pAldA | *PSPTO_0092* CDS and promoter region in pME6031; Tet^r^ | (McClerklin et al., 2018) |
| pJP5603-*aldA*::Ω | *aldA* marker replacement plasmid in pJP5603; Spec^r^, Km^r^ | This study |
| pHP45 | plasmid with the omega fragment containing spec resistance, Spec^r^ | (Prentki and Krisch, 1984) |
| pJP5603-2673int | Insertion/disruption plasmid for *aldB*; Km^r^ | (McClerklin et al., 2018) |
| pET28a-AldA | HIS-tagged AldA expression construct; Km^r^ |  |

**Table S2.** Primers used in this study.

| **Name** | **Sequence (5’ to 3’)** |
| --- | --- |
| **For cloning** |  |
| aldA_up_F | AGGAAACAGCTATGACCATGATGTCGAGAGTATGTCAAG |
| aldA_up_R | TCACCGGATCTTTTATGTCCTCCTCTTGTTG |
| omega_frag_F | GGACATAAAAGATCCGGTGATTGATTGAG |
| omega_frag_R | AGGTTGCGCCCGGTGATTGATTGAGCAAG |
| aldA_down_F | TCAATCACCGGGCGCAACCTTTACGCTTG |
| aldA_down_R | GTTGTAAAACGACGGCCAGTAAAGCGGCAACAGGGTGAG |
| pJP5603_F | ACTGGCCGTCGTTTTACAAC |
| pJP5603_R | CATGGTCATAGCTGTTTCCTGTGTGAAATTG |
| **For genotyping** |  |
| omega_out_up | CTGGAAGGCGAGCATCGTTTG |
| omega_out_down | ACTCAAGCGTTAGATGCACTAAGC |
| M13F | GTAAAACGACGGCCAG |
| M13R | CAGGAAACAGCTATGACC |
| PSPTO_0089_ups_F | TTCACTGTAGGTGTAGCCGGTAAG |
| PSPTO_0094_R | GATCAGCATCAGCAGGCCAAAGG |
| pME6031_for | ATCCTTGACCCGCAGTTGCAAACCC |
| Shuttle_rev | AAGGTTATCAAGTGAGAAATCACCA |
| 0092seqF | CGTACTGGTTGACCCACA |
| 0092seqR | GAACAACGCGCCCAAAATC |
| Kan2 | AGCCGAATAGCCTCTCCA |
| 2673SeqF | ATCCCTGAACGAATGTCCCG |

**Table S3.** Metabolites detected by LC-MS/MS**.**

| Metabolite^a^ | Precursor ion (*m*/*z*) | Product ion (*m*/*z*) | Retention time (min) | Fragmentor  (V) | Collision energy  (V) |
| --- | --- | --- | --- | --- | --- |
| IAA | 176.1 | 130.1 | 6.0 | 60 | 13 |
| IAAld | 160.1 | 118.1 | 5.9 | 110 | 9 |
| IAA-Lys | 304.2 | 130.1 | 5.2 | 109 | 44 |
| IAM | 175.1 | 130.1 | 5.4 | 35 | 13 |
| TAM | 161.1 | 144.1 | 4.3 | 60 | 9 |
| PAA | 137.1 | 91.1 | 5.6 | 60 | 13 |
| PAAld | 121.1 | 79.1 | 1.5 | 60 | 5 |
| Trp | 205.2 | 188.2 | 4.0 | 60 | 5 |
| Phe | 166.2 | 120.1 | 1.2 | 76 | 9 |

^a^IAA, Indole-3-acetic acid; IAAld, indole-3-acetaldehyde; IAA-Lys, IAA-lysine; IAM, indole-3-acetamide; TAM, tryptamine; PAA, phenylacetic acid; PAAld, phenylacetaldehyde; Trp, tryptophan, Phe, phenylalanine.

**Table S4.** Accumulation of phenylacetic acid (PAA) in conditioned medium in three independent experiments.

| [PAA µM]^a^ in media | | | |
| --- | --- | --- | --- |
| Media Expt. date: | 3/2/2019 | 11/2/2018 | 4/11/2018 |
| HSC | n.d. | n.d. | 0.011 |
| HSC + PAAld | n/a | 0.284 | 0.513 |
| cond. HSC^b^ (WT) | n.d. | 0.359 | n/a |
| cond. HSC (WT) + PAAld | 23.379 | 21.129 | n/a |
| cond. HSC (*aldA*) | n.d. | n/a | n/a |
| cond. HSC (*aldA*) + PAAld | 32.809 | n/a | n/a |

^a^ PAA levels were measured using LC-MS/MS 46-48 hrs after incubating HSC supplemented with 25 µM phenylacetylaldehyde (PAAld). No bacterial cells were included.

^b^ Conditioned HSC was made by growing *Pto*DC3000 in HSC for 48 hrs, removing the cells by centrifugation, followed by filtering the supernatant with a 0.2-micron filter.

n.d., not detected; n/a, not analyzed.
